# Contactin-1 is a critical neuronal cell surface receptor for perineuronal net structure

**DOI:** 10.1101/2024.11.05.622114

**Authors:** Ashis Sinha, Gabrielle Nickerson, Samuel Bouyain, Russell T. Matthews

## Abstract

Perineuronal nets (PNNs), are neuron-specific substructures within the neural extracellular matrix (ECM). These reticular structures form on a very small subset of neurons in the central nervous system (CNS) and yet have a profound impact in regulating neuronal development and physiology. PNNs are well-established as key regulators of plasticity in the CNS. Their appearance coincides with the developmental transition of the brain more to less plastic state. And, importantly, numerous studies have demonstrated that indeed PNNs play a primary role in regulating this transition. There is, however, a growing literature implicating PNNs in numerous roles in neural physiology beyond their role in regulating developmental plasticity. Accordingly, numerous studies have shown PNNs are altered in a variety of neurological and neuropsychiatric diseases, linking them to these conditions. Despite the growing interest in PNNs, the mechanisms by which they modulate neural functions are poorly understood. We believe the limited mechanistic understanding of PNNs is derived from the fact that there are limited models, tools or techniques that specifically target PNNs in a cell-autonomous manner and without also disrupting the surrounding neural ECM. These limitations are primarily due to our incomplete understanding of PNN composition and structure. In particular, there is little understanding of the neuronal cell surface receptors that nucleate these structures on subset of neurons on which they form in the CNS. Therefore, the main focus our work is to identify the neuronal cell surface proteins critical for PNN formation and structure. In our previous studies we demonstrated PNN components are immobilized on the neuronal surface by two distinct mechanisms, one dependent on the hyaluronan backbone of PNNs and the other mediated by a complex formed by receptor protein tyrosine phosphatase zeta (RPTPζ) and tenascin-R (Tnr). Here we first demonstrate that the Tnr-RPTPζ complex in PNNs is bound to the cell surface by a glycosylphosphatidylinositol (GPI)-linked receptor protein. Using a biochemical and structural approach we demonstrate the GPI-linked protein critical for binding the Tnr-RPTPζ complex in PNNs is contactin-1 (Cntn1). We further show the binding of this complex in PNNs by Cntn1 is critical for PNN structure. We believe identification of CNTN1 as a key cell-surface protein for PNN structure is a very significant step forward in our understanding of PNN formation and structure and will offer new strategies and targets to manipulate PNNs and better understand their function.

## INTRODUCTION

Perineuronal nets (PNNs) are a mesh-like conspicuous sub-compartment of the neural extracellular matrix (ECM) occurring around the cell bodies and proximal dendrites of subsets of neurons in the central nervous system (CNS). PNNs are assembled as stable, mature synaptic circuitry being established in the CNS, which closes the highly plastic critical period (CP) in the animal’s neuronal development. A major driver behind the formation of nets is neuronal activity and, accordingly, decreased neuronal firing during early postnatal life leads to attenuated formation of PNNs. For example, rearing animals in complete darkness delays PNN formation and CP closure in the primary visual cortex and depriving sensory input by whisker trimming before CP closure leads to decreased PNN expression in the barrel cortex (1–4). Interestingly, if these procedures are carried out after CP closure no effects on PNNs are observed. The temporal appearance of PNNs during development and their activity-dependent formation has led investigators to postulate their involvement in the stabilization and maturation of synapses during development and in restricting plasticity in adult animals (1, 3, 5, 6). PNNs are particularly enriched in the glycosaminoglycan hyaluronan (HA) and hyaluronan-binding chondroitin sulfate proteoglycans (CSPGs). Digesting away PNNs by injecting bacterial chondroitinase ABC (ChABC), an enzyme capable of degrading the HA backbone of PNNs and the chondroitin sulfate chains of CSPGs, Pizzorusso *et al*. were able to reopen juvenile forms of plasticity in the visual cortex of adult rats (3). In addition to the cortex, a large amount of work has also demonstrated the role of PNNs in regulating plasticity across various other brain regions including the amygdala, hippocampus, striatum, and cerebellum (7–12). However, despite this increasing amount of evidence demonstrating the role of PNNs in regulating plasticity in the CNS, a mechanistic understanding of PNN function has been elusive.

The limited grasp of PNN function is derived primarily from our incomplete understanding of their structure and molecular composition and in turn, our inability to disrupt them specifically without also disrupting the surrounding ECM. Identifying the role of PNNs has relied primarily on enzymatic manipulation of these structures using ChABC. Though treatment with ChABC affects PNNs, it does not necessarily eliminate them and broadly disrupts any HA based ECM structure in the local area making it difficult to clearly delineate out the role of these enigmatic structures in the CNS. Additionally, almost all known PNN components are secreted molecules and are broadly expressed in the neural ECM (13, 14). PNNs however, are distinct from the surrounding ECM and form around only subsets of cells, specifically interneurons expressing the Ca^2+^-binding protein parvalbumin. Genetic models targeting specific PNN components exist and have provided valuable insight into PNN function. However, to date they have focused on broadly secreted PNN components and thus, a mechanism of PNN specificity has remained elusive (14–23). Therefore, the goal of our study is to provide a more complete understanding of PNN formation and structure. Critically, this includes identifying the cell-surface receptors that are critical for PNNs.

In our previous studies we demonstrated that receptor protein tyrosine phosphatase zeta (RPTPζ) is essential for PNN formation, as is its binding to the extracellular matrix glycoprotein tenascin-R (Tnr) (22, 24). Along with other existing data on PNN structure, our findings show that PNN components are immobilized on the neuronal surface by two distinct mechanisms. One is mediated by Tnr/RPTPζ complex while the other appears dependent on the HA backbone of PNNs. Both RPTPζ and Tnr are necessary for the lattice or net-like appearance of PNNs and the absence of either leads to disrupted and aggregated PNN structures. RPTPζ exists in multiple secreted and membrane-bound forms created by alternative splicing as well as proteolytic cleavage (25–27). Among these, however, we found that the secreted splice variant of RPTPζ called phosphacan is the critical form in PNNs, which left the question of how the Tnr/RPTPζ complex immobilizes PNNs to the cell surface unanswered. In this present study we identify the receptor for this complex in PNNs as the glycosylphosphatidylinositol (GPI)-linked protein, contactin-1 (Cntn1). Knockout of *Cntn1* is sufficient to alter drastically PNN through disrupted binding of the Tnr/RPTPζ complex. To our knowledge this is the first direct evidence uncovering the role of Cntn1 in PNN formation and the first identification of a cell surface receptor that is critical for PNN structure. The findings presented here provide novel insights into the structure and formation of PNNs and offer new strategies to manipulate them and better understand their function.

## RESULTS

### Disrupted PNN staining in Ptprz1 KO neuronal cultures is rescued by purified or recombinant phosphacan

As we demonstrated in our previous work, PNN structure is disrupted in *Ptprz1* KO mice which carry null alleles for the CSPG RPTPζ (22, 24). In cultured neurons from these mice, the disrupted PNN structure can be visualized by staining for the key PNN component aggrecan (21, 28, 29). In wildtype (WT) neuronal cultures aggrecan staining presents with a lattice-like pattern on the cell surface but is discontinuous and punctate in *Ptprz1* KO cultures (Fig. 1). RPTPζ exists in multiple isoforms including transmembrane and secreted forms (25–27). But consistent with our previous studies, PNN structures on *Ptprz1* KO neurons are rescued by adding a soluble form of RPTPζ known as phosphacan (Fig. 1, Eill *et al*. (22)). And, importantly, PNN structures are similarly rescued using a recombinant form of RPTPζ that lacks the middle glycosaminoglycan attachment region (Fig. 1, Sinha *et al*. (24)). These data confirm that a secreted form of RPTPζ is critical for PNN structure and that the essential interactions mediated by RPTPζ are independent of its chondroitin sulfate chains and likely mediated by protein-protein interactions. However, it also suggests that since it is the secreted form of RPTPζ, phosphacan, that mediates these effects on PNNs, that there must be additional proteins and potentially a cell surface receptor that link phosphacan to the neuronal surface in PNNs.

**Figure 1.**
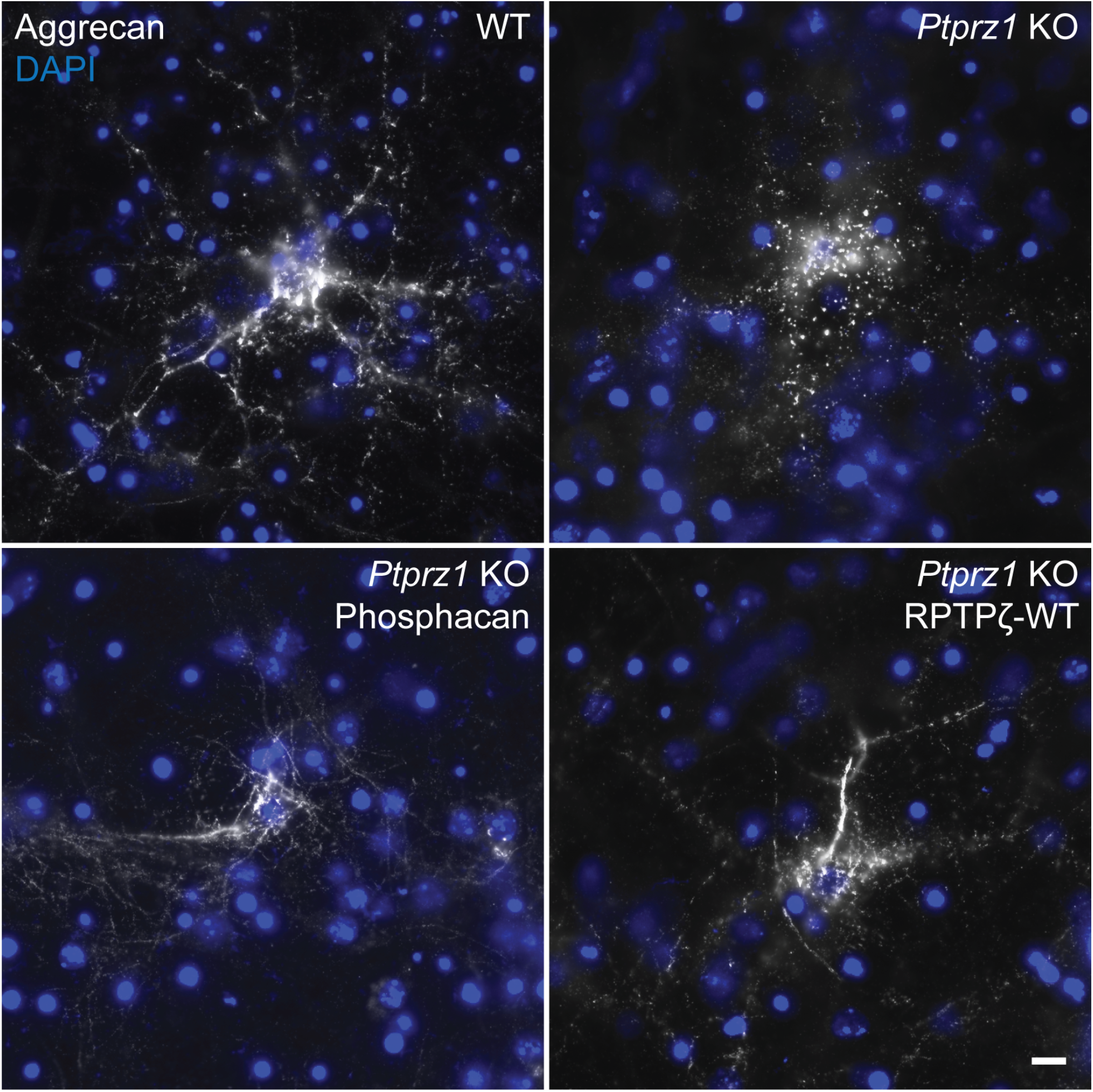
Disrupted PNN Structure in *Ptprz1* KO cultures can be recovered with purified phosphacan or recombinant RPTPζ. Neurons derived from E16 WT mice were positive for PNN component aggrecan and displayed regular and continuous staining. Aggrecan staining in cultures from *Ptprz1* KO mice was disrupted and appeared broken, aggregated, and punctate. Addition of phosphacan purified from WT brain homogenates to *Ptprz1* KO cells restored PNN component binding and structure. As above, addition of recombinant RPTPζ (RPTPζ-WT) to *Ptprz1* KO cells restored PNN component binding and structure. These findings are consistent with previously published studies(22, 24). Scale bar, 10 μm.

### Membrane binding of key PNN component aggrecan is partially dependent on a glycophosphatidylinositol (GPI)-anchored protein

In our previous work we developed a testable model of PNN structure. In this model we hypothesized a key PNN component, aggrecan, is bound to the HA backbone of PNNs by its N-terminus and to Tnr by its C-terminus in a Ca^2+^ -dependent manner (30–32). Tnr is then bound to phosphacan. Consistent with this model, our previously published work has demonstrated that protein-protein interactions between Tnr and phosphacan are essential for PNN structure. Phosphacan presumably would then be bound to the cell surface by a cell-surface protein. However, phosphacan is a known binding partner of several cell adhesion molecules such as Cntn1, Ng-CAM, N-CAM and Nr-CAM (33–37). These cell surface receptors include both transmembrane proteins and GPI-anchored proteins. In order to narrow down the list of potential cell-surface binding partner for phosphacan in PNNs, we turned to a biochemical release assay we have developed in the lab(22). Using membranes isolated from adult mouse brain we determined that aggrecan, the most specific PNN component, is bound to the neuronal cell surface in PNNs by two distinct mechanisms, one dependent on the HA backbone, susceptible to enzymatic digestion of HA, and the other dependent on Ca^2+^ ions, susceptible to chelation by EDTA/EGTA (22). Importantly, we previously found that in membranes from *Ptprz1* KO brains, aggrecan binding became solely dependent on HA and was no longer also bound in a Ca^2+^-dependent manner (22). We reasoned if phosphacan was bound via a GPI-linked protein then cleaving the GPI-linkage with phosphatidylinositol-specific phospholipase-C (PIPLC) should be able to replace Ca^2+^ chelation in the release of aggrecan. If treatment with PIPLC did not lead to the release of aggrecan, then we could rule out GPI-anchored proteins from being involved in binding phosphacan integrated into PNNs.

Membrane fractions from post-natal day 45 (PND 45) *Ptprz1* WT mouse brains were isolated and treated with ChABC to digest the HA backbone of PNNs, and/or EDTA to chelate Ca^2+^ ions, and/or PIPLC to disrupt any GPI links. After treatment, samples were centrifuged to obtain soluble release (R) and insoluble pellet (P) fractions and the release of aggrecan was analyzed by Western blot (Fig. 2A). We found significant differences in the release of aggrecan among the various treatment groups (one-way ANOVA F (6, 16) = 61.31, *p* < 0.0001) (Fig. 2B). As expected, aggrecan was partially released by ChABC treatment alone (54 ± 5%, *p* < 0.0001). However, a combination of ChABC and EDTA led to an almost complete release of aggrecan into the soluble phase (79 ± 7%, p < 0.0001). Interestingly, we also found that the majority of aggrecan is released into the soluble phase by ChABC and PIPLC treatment (86 ± 6%, *p* < 0.0001). Release of aggrecan by both ChABC and EDTA (*p* < 0.0001) and the ChABC and PIPLC treatment (*p* < 0.0001) was significantly higher than by ChABC treatment alone suggesting that in addition to the HA backbone of PNNs, aggrecan is bound to the cell membrane by Ca^2+^ dependent mechanism and a GPI anchor-dependent protein. EDTA (30 ± 3%, *p* = 0.0141) and PIPLC (44 ± 3%, *p* < 0.0001) treatment alone had significant effects on aggrecan release as well, indicating the role of these interactions in binding aggrecan to the cell surface.

**Figure 2.**
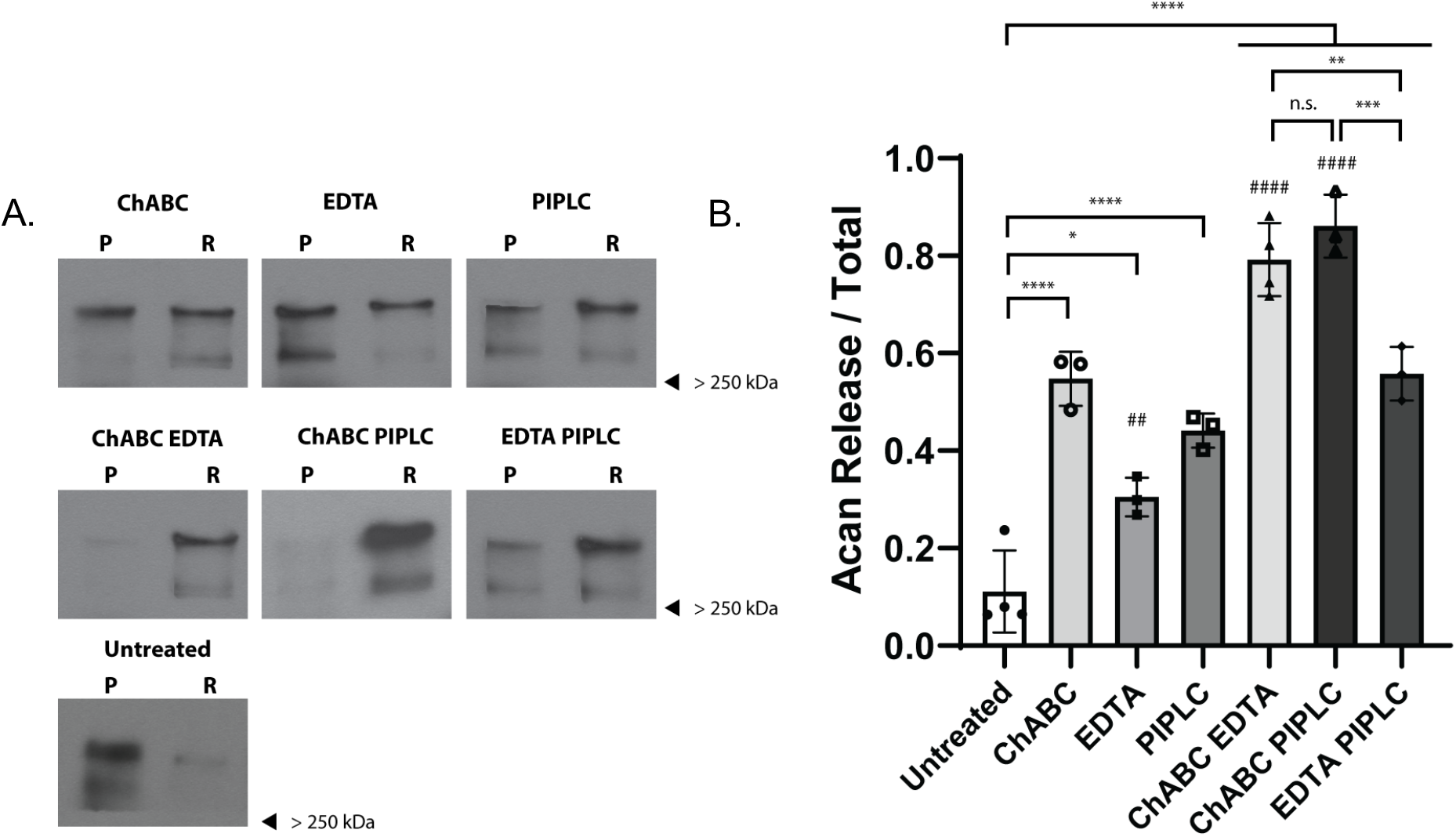
Membrane binding of key PNN component aggrecan depends on the HA backbone, Ca^2+^ ions and a GPI-anchored protein. Homogenates PND 45 WT brains were treated with ChABC to digest the hyaluronan backbone of PNNs and/or EDTA to chelate Ca^2+^ ions and/or PIPLC to disrupt any GPI linked proteins. After treatment samples were centrifuged to obtain insoluble pellet (P) and soluble release (R) fractions. *(A)* Western blotting image showing release of PNN marker aggrecan into soluble phase from brain homogenates of WT mice by ChABC, EDTA and PIPLC treatment alone and combinations of ChABC EDTA, ChABC PIPLC and EDTA PIPLC treatment. *(B)* There was a statistically significant difference in release of aggrecan across various treatment groups as determined by one-way ANOVA (F (6, 16) = 61.31, *p* < 0.0001). Aggrecan is partially released into the soluble fraction by ChABC (54 ± 5%, *p* < 0.0001), EDTA (30 ± 3%, *p* = 0.0141) and PIPLC (44 ± 3%, *p* = 0.0001) treatment alone as compared to untreated condition. However, aggrecan is almost completely released into the soluble phase by ChABC EDTA (79 ± 7%, *p* < 0.0001) and ChABC PIPLC treatment (86 ± 6%, *p* < 0.0001) as compared to the untreated group. Release of aggrecan by ChABC EDTA (*p* < 0.0001) and ChABC PIPLC treatment (*p* < 0.0001) was significantly greater than by ChABC treatment alone. Combination of EDTA PIPLC led to a significantly greater release of aggrecan than untreated group (55 ± 5%, p < 0.0001). However, it was not significantly different from treatment with ChABC, EDTA or PIPLC alone. * in *(B)* indicates significance compared to untreated group unless denoted otherwise, # indicates comparison to ChABC treated group. Bar in graphs represent percentage release ± S.D.

We reasoned that if treatments with EDTA and PIPLC affected two distinct anchoring mechanisms, then the release of aggrecan with a combination of Ca^2+^ chelation and ablation of GPI anchors should be more significant than for each group alone. Combining these two treatments did result in a significant release of aggrecan as compared to the untreated group (55 ± 5%, p < 0.0001). However, the release was not significantly greater than in each of the individual treatment conditions. Furthermore, treatments with EDTA, PIPLC or a combination of the two were not significantly different from treatment with ChABC alone. These findings are consistent with the hypothesis that PNNs are immobilized to the cell membrane by two distinct mechanisms, one involving HA and the other involving Ca^2+^. Results from these aggrecan-release experiments also indicate to us that a GPI-linked protein is critically involved in linking aggrecan to the cell membrane. We speculate that this GPI-anchored protein is part of the Ca^2+^-dependent release mechanism of aggrecan and thus is implicated in the binding of phosphacan to the cell surface.

### PNN components are immobilized on the cell surface, in part, by a GPI-linked protein

The Western blot analyses described above made us speculate that phosphacan is likely the PNN component that is bound via a GPI-linked protein and led us to form the model structure shown in Figure 3A. An interesting outcome of this model is that it predicts aggrecan would be released from PNN by HA digestion combined with either Ca^2+^ chelation or PIPLC digestion but that other components such as Tnr would likely only be released by PIPLC digestion and not by Ca^2+^ chelation. However, unlike aggrecan, Tnr is found both in PNNs and throughout the neural ECM so that we decided to test this hypothesis by analyzing the effects of ChABC, EDTA, and PIPLC treatments on PNN structure in cultured primary neurons.

**Figure 3.**
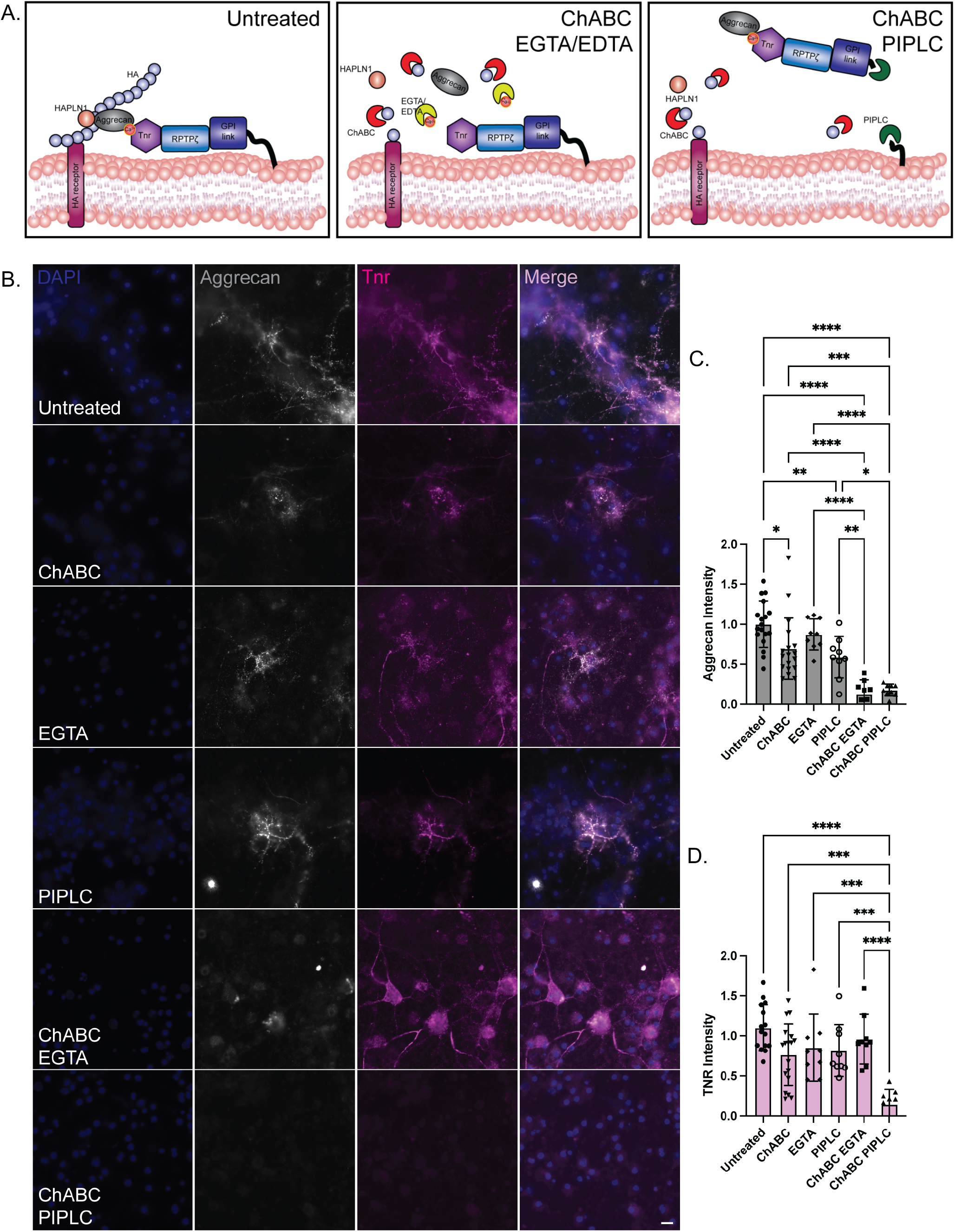
Treatment with ChABC and PIPLC eliminates staining for PNN components aggrecan and Tnr. *(A)* Based on our initial studies and work done previously we created a model of PNN structure including a GPI-linked protein that tethers the structure to the cell surface. We hypothesize that aggrecan binds to Tnr, which is then bound to RPTPζ/phosphacan, and the entire complex is attached to the cell surface by a membrane receptor. The interaction of aggrecan and Tnr is known to be Ca^2+^ dependent, therefore, aggrecan is released by a combination of HA digestion and Ca^2+^ chelation, but Tnr bound to phosphacan and subsequently a membrane receptor (possibly a GPI-anchored protein) remains tethered to the cell membrane. We predict that combining HA digestion with PI lipid specific phospholipase-C (PIPLC) treatment will release both aggrecan and Tnr from the cell surface. *(B)* To test this model, we turned to neuronal cell culture. Neurons derived from E16 WT mice were positive for PNN components aggrecan and Tnr. Acute treatment with ChABC, EGTA or PIPLC alone only had small effects on aggrecan and Tnr levels. Treatment with ChABC to digest away the HA backbone of PNNs and EGTA to chelate Ca^2+^ ions in combination eliminated aggrecan staining but only had a minor effect on Tnr levels. Treatment with ChABC and PIPLC eliminated both aggrecan and Tnr staining. (*C, D*) To quantify these findings, aggrecan and Tnr intensity for the different treatment groups relative to untreated condition was calculated and is presented here in a graphical form. Analysis by one-way ANOVA showed significant differences in levels of aggrecan and Tnr among different treatment groups (aggrecan F (5, 64) = 29.12, *p* < 0.0001; Tnr F (5, 63) = 9.92, *p* < 0.0001). Tukey’s *post hoc* testing showed significant loss of aggrecan but not Tnr with ChABC and EGTA treatment (aggrecan *p* < 0.0001; Tnr *p* = 0.9123). There was significant loss of both aggrecan and Tnr with ChABC and PIPLC treatment (aggrecan *p* < 0.0001; Tnr *p* < 0.0001). Tnr release was significantly greater with ChABC and PIPLC compared to ChABC and EGTA (p < 0.0001). These results indicate that aggrecan is immobilized on the cell surface by a HA dependent mechanism then by a Ca^2+^ sensitive interaction and further downstream by a GPI anchor. Tnr on the other hand acts as an adapter molecule between aggrecan and the GPI anchor. Tnr is bound to the HA backbone through its interaction with aggrecan and remains attached to the cell surface by the GPI anchor when treated with ChABC and EGTA. Tnr staining is lost only when both the HA backbone and GPI anchor is disrupted by ChABC and PIPLC treatment. Bar in graphs represent percentage release ± S.D. Scale bar, 10 μm. * in *(C & D)* indicates significance.

Dissociated neuronal cultures present an attractive model to study interactions required for the formation of PNNs as they allow us to carry out our manipulations in live cells. Using this model system we have previously shown that aggrecan staining in PNNs is profoundly disrupted by the combination of digestion of HA and chelation of Ca^2+^ but only slightly impacted with either treatment alone (22). Here we additionally investigated if PIPLC could replace Ca^2+^ chelation in disrupting PNN structures. We found that treating cultured neurons with either ChABC, or EGTA only had small effects on aggrecan reactivity in PNNs (Fig 3B and 3C.). Moreover, aggrecan levels under these treatment conditions were not significantly different from untreated controls. However, a combination of either ChABC and EGTA (loss = 87 ± 17%, *p* < 0.0001, Tukey’s *post hoc* testing) or ChABC and PIPLC (loss = 82 ± 7%, *p* > 0.0001, Tukey’s *post hoc* testing) led to an almost complete elimination of aggrecan staining.

Intriguingly when we expanded these analyses to another essential PNN component, Tnr, the results provided greater insight into PNN structure (Fig 3B & 3D) and remained consistent with our model (Fig 3A). The combination of ChABC and EGTA had no significant impact on Tnr levels (loss = 13 ± 14%, *p* > 0.999, Tukey’s *post hoc* testing), whereas treating with ChABC and PIPLC essentially eliminated Tnr staining (loss = 85 ± 18%, *p* < 0.0001, Tukey’s *post hoc* testing). As mentioned above we hypothesized aggrecan is bound to the HA backbone of PNNs by its N-terminus and to Tnr by its C-terminus in a Ca^2+^ dependent manner. Tnr is then bound to phosphacan, and this complex is retained at the cell surface by a membrane receptor (Fig. 3A). Therefore, whereas aggrecan can be released by a combination of HA digestion and Ca^2+^ chelation a complex of Tnr and phosphacan cannot because it remains tethered to the cell surface by a GPI-anchored protein. Combining HA digestion with PIPLC treatment releases both aggrecan and Tnr from the cell surface. Importantly, these results show that the protein tethering PNNs to the cell surface is a GPI-linked protein.

### PNN structure can be disrupted by functionally blocking Cntn1

Our studies above indicated a GPI-linked protein is critically involved in PNN structure and that treatment with PIPLC can replace Ca^2+^ chelation to disrupt PNN components. In our previous work we showed that in *Ptprz1* KOs, PNNs are no longer bound to the neuronal surface in a Ca^2+^ dependent manner and that Tnr and phosphacan form a key complex in PNNs (22, 24). Together these findings led us to believe that phosphacan binding in PNNs is dependent on a GPI-linked protein. To the best of our knowledge, the only GPI-linked protein that is known to interact with phosphacan is Cntn1 (36, 37). Interactions between RPTPζ/phosphacan and Cntn1 are essential to the maturation of oligodendrocyte precursor cells and are also known to promote the outgrowth of neurites in cultured neurons (35–37). Cntn1, however, has never previously been associated with PNNs.

As a first step to determine if Cntn1 might contribute to PNN structure we utilized our primary neuronal culture model to determine if a function blocking antibody directed against CNTN1 could disrupt PNNs. As a control we used an antibody directed against Cntn4. Importantly Cntn1 interacts specifically with phosphacan and does not bind with other members of the family, Cntn2-6 (36). As described above, we derived cultures from embryonic day 16 (E16) CD1 WT mice embryos and to block the interaction of any protein with Cntn1, we added an anti-Cntn1 polyclonal antibody (goat IgG, final concentration of 2.5 μg/ml) or the same concentration of an anti-Cntn4 antibody to the cultures at 6 days *in vitro* (DIV 6) (38). We fixed the cells at DIV 9 and analyzed the binding of PNN component aggrecan with PNN peak/node analysis (22, 24). We identified aggrecan nodes or peaks using the local maxima function in ImageJ and determined the difference in intensity between the nodes and their surrounding space on the cell surface (mean node prominence) using an *ad hoc* algorithm. We found that untreated cells were positive for the PNN marker aggrecan and displayed regular PNN structures (Fig. 4). PNN staining in control cells (Untreated and anti-Cntn4 antibody addition) appeared continuous with PNN peaks or nodes connected by bridge-like internode staining. In contrast, cells treated with the Cntn1 neutralizing antibody showed disrupted PNN structures (Fig. 4). PNN peaks in cultures where interaction with Cntn1 was blocked were significantly more prominent and isolated. Aggrecan staining appeared discontinuous and aggregated and resembled PNNs from *Ptprz1* KO cultures (Fig. 1) (one-way ANOVA F(2, 46) = 7.63., *p* < 0.0014, mean node prominence untreated = 5.55 ± 1.00, anti-Cntn1 antibody = 7.59 ± 2.45, p = 0.0023, anti-Cntn4 antibody = 5.71 ± 1.41, p = 0.95). Moreover, mean node prominence between anti-Cntn1 and anti-Cntn4 antibodies showed statistically significant differences (p = 0.0078). These results indicate that Cntn1 is required for proper binding of PNN components to the neuronal surface and is responsible for the regular lattice formation seen in untreated cells. Furthermore, these experiments demonstrate that binding of PNN components can be disrupted by blocking their interactions with Cntn1.

**Figure 4.**
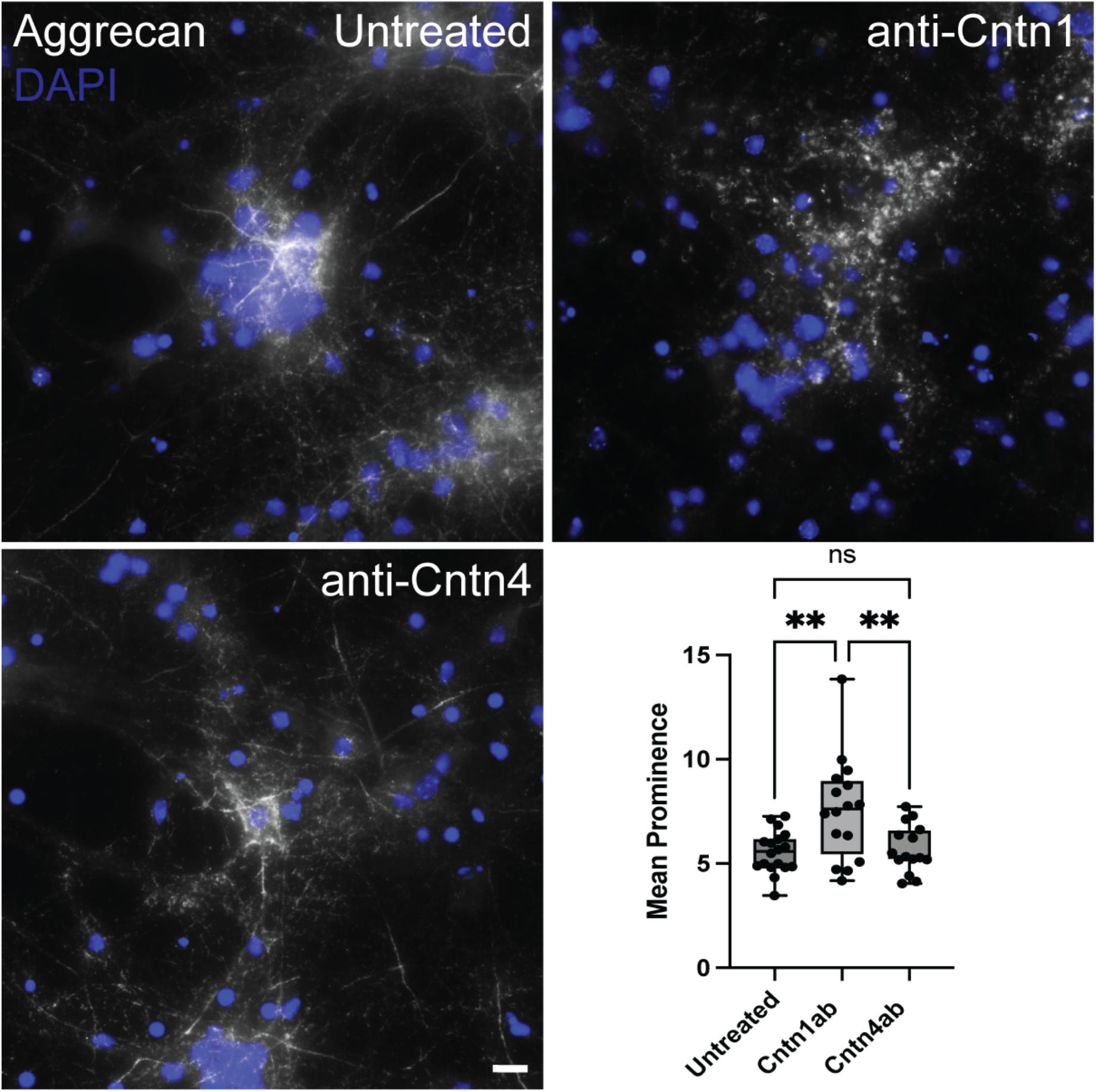
PNNs structure is disrupted when Cntn1 is functionally blocked in culture. Cortical cultures were derived from E16 WT mice. Cells were fixed at DIV 9 by and PNNs were visualized by staining with PNN marker, aggrecan. PNNs staining appeared regular and continuous in untreated group. Addition of anti-Cntn1 function-blocking antibody (2.5 μg at DIV 6) resulted in disrupted and aggregated PNN structures and isolated PNN node/peaks. PNN structure was not disrupted by addition of anti-Cntn4 antibody (2.5 μg at DIV 6). Mean prominence or isolation of PNN peaks was quantified using our PNN node/ peak analysis. Analysis by one-way ANOVA showed significant differences in the mean prominence index of PNNs (determined by the average isolation index of aggrecan staining peaks) among different treatment groups (aggrecan F (2, 46) = 7.63, *p* = 0.0014). Tukey’s *post hoc* testing showed significant difference in mean prominence of aggrecan peaks between untreated and anti-Cntn1 antibody treatment group (p = 0.0023) and between anti-Cntn1and anti-Cntn4 treatment (p = 0.0078). No significant difference was observed between Untreated and anti-Cntn4 antibody treatment group (p = 0.9582). n = 15-20 PNNs per treatment group, 3-4 independent cultures. Scale bar, 10 μm.

### *PNN structure depends on* RPTPζ *binding to* Cntn1

As we have showed in previous studies and above (Fig. 1), PNNs are disrupted in cultured neurons derived from *Ptprz1* KOs (22). However, PNN structure can be restored by addition of brain-purified phosphacan or by addition of a recombinant truncated phosphacan construct. Importantly, previous structural work also provided atomic-level insights into the interaction between phosphacan and Cntn1 (36, 37). These studies demonstrated that a β-hairpin loop in the N-terminal carbonic anhydrase-like domain of phosphacan is critical for interacting with Cntn1. Using this information, the authors created a mutant form of phosphacan in which nine amino acids are deleted from this loop and that no longer interacts with Cntn1 (named here RPTPζ-βdel). Here we compared the ability of the RPTPζ-βdel protein to recover PNN structure in cultured neurons compared to the wildtype RPTPζ protein (RPTPζ-WT). As described in neuronal cultures derived from E16 *Ptprz1* KO mice pups show irregular and aggregated staining for the PNN component aggrecan when fixed at 9 DIV (Fig. 5A). As expected RPTPζ-WT protein added exogenously at 3 DIV recovered and restored aggrecan binding to the cell surface in *Ptprz1* KO neurons. In contrast, the Cntn1-binding mutant, RPTPζ-βdel, did not restore the PNN aggrecan structure in cultured neurons. To quantitatively assess this structural recovery, we utilized PNN peak/node analysis (Fig 5B). We found statistically significant differences in mean PNN node prominence in our various treatment groups (one-way ANOVA F (2, 52) = 6.98, *p* = 0.0021). Addition of RPTPζ-WT recovered PNN structure in *Ptprz1* KO cells (mean node prominence: Untreated = 26 ± 7.29, RPTPζ-WT 19.55 ± 3.19, p = 0.0017, Tukey’s *post hoc* testing). In contrast, RPTPζ-βdel, did not restore the PNN aggrecan structure in *Ptprz1* KO cultured neurons and aggrecan staining on cells continued to appear discontinuous and aggregated (mean node prominence = 23.91 ± 5.22, p = 0.4352). Moreover, mean node prominence between RPTPζ-WT and RPTPζ-βdel treatment groups also showed statistically significant differences (p = 0.0429). Together these data show that the binding of phosphacan to Cntn1 is critical for PNN structure.

**Figure 5.**
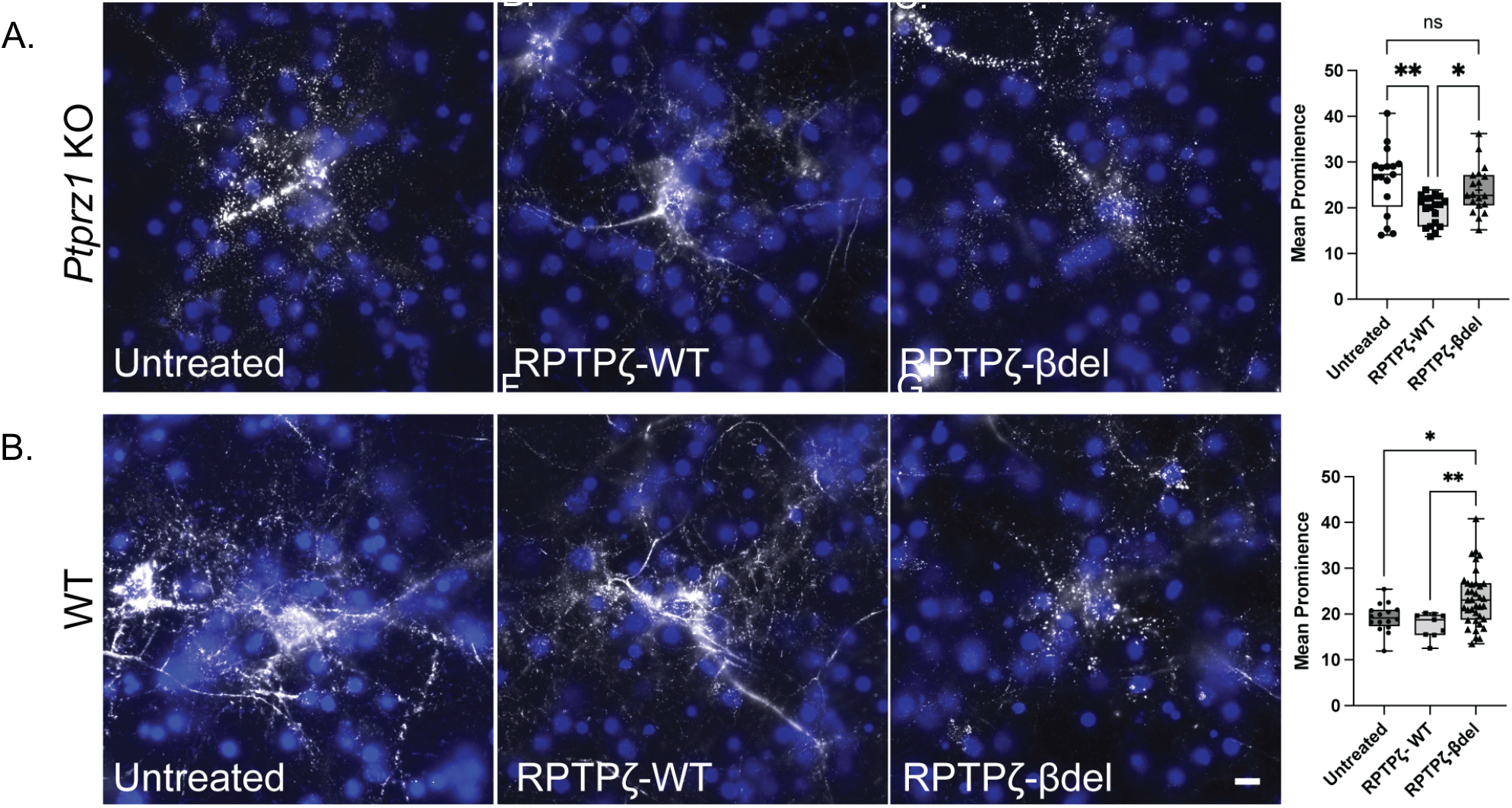
Addition of RPTPζ mutants is unable to recover PNN structure in RPTPζ deficient *Ptprz1* KO neuronal cultures and disrupts PNNs in WT cultures. Cortical cultures from E16 *Ptprz1* KO mice lacking RPTPζ (A) or WT embryos (B) were derived and fixed at DIV 9. (A) Binding of PNN component aggrecan appeared disrupted with a broken and aggregated staining pattern in *Ptprz1* KO cultures. Addition of recombinant phosphacan RPTPζ-WT (2 μg per well at DIV 3) to cells was able to restore aggrecan binding. Addition of Cntn1 binding mutant RPTPζ-βdel was unable to restore PNNs and aggrecan staining continued to appear broken and aggregated. Analysis by ordinary one-way ANOVA showed average isolation index of aggrecan staining nodes representing mean prominence of PNN staining peaks is significantly different among groups (F (2,52) = 6.98, *p* = 0.0021). RPTPζ-WT treated cells (n = 19 cells, 3 cultures) showed a significantly lower mean prominence (*p* = 0.0017) compared to untreated cells (n = 17 cells, 3 cultures) indicating more regular pattern of aggrecan staining and a decrease in node/ peak isolation. Mean prominence of peaks in RPTPζ-βdel treated PNNs was not significantly different (n = 19 cells, 4 cultures, *p* = 0.4352) from untreated cells. Mean prominence of peaks in RPTPζ-βdel group was also significantly higher than RPTPζ-WT treated cells (p = 0.0429). (B) Cortical cultures derived from E16 WT mice fixed at DIV 9 and stained with PNN marker, aggrecan *s*howed regular and continuous aggrecan staining in untreated group. Addition of RPTPζ-WT did not have any significant effects on aggrecan staining. Addition of RPTPζ-βdel (2 μg per well at DIV 6) resulted in disrupted and aggregated aggrecan staining. Analysis by ordinary one-way ANOVA showed significant differences in the mean prominence of PNNs (average isolation index of aggrecan staining peaks) among the various treatment groups (F (2, 59) = 6.91, *p* = 0.0020). Mean prominence is not significantly higher in RPTPζ-WT treated cells (n = 9 PNNs, 3 cultures, *p* = 0.7201) as compared to untreated control cells (n = 15 cells, 4 cultures). On the other hand, mean prominence is significantly higher in RPTPζ-βdel treated cells (n = 37 PNNs, 7 cultures, *p* = 0.0208) as compared to untreated control cells. Scale bar, 10 μm.

Having determined that binding to Cntn1 is critical to recover PNN structure in *Ptprz1* KO neurons, we next asked if RPTPζ-βdel would act as a dominant negative and disrupt PNN structure in wildtype neurons (Fig. 5B). To quantitatively assess changes in PNN structure, we again utilized our PNN peak/ node analysis (Fig 5B). Here also, we found statistically significant differences in mean PNN node prominence among our various treatment groups (one-way ANOVA F (2, 59) = 6.915, *p* = 0.0020). In untreated wildtype neurons, aggrecan staining appeared normal (mean node prominence: Untreated = 19.19 ± 3.10), and this was unaltered by the addition of RPTPζ-WT (mean node prominence: RPTPζ-WT = 17.56 ± 2.71, p = 0.8632). In contrast, addition of RPTPζ-βdel disrupted aggrecan staining and made it appear discontinuous and aggregated in appearance, similar to the aggrecan staining seen in *Ptprz1* KO neurons (mean node prominence: RPTPζ-βdel = 23.38 ± 6.03, p = 0.0003). Mean node prominence in RPTPζ-βdel treatment group was also significantly higher than RPTPζ-WT group (p = 0.0085). Together these data demonstrate that binding of phosphacan to Cntn1 is necessary for PNN structure.

### PNN structure is disrupted in mice carrying null alleles for Cntn1

Our results from *in vitro* studies suggest that PNN components are immobilized to the neuronal surface by interacting with GPI-linked protein, Cntn1. If this is indeed true, then PNNs should appear disrupted in mice carrying null alleles for *Cntn1*. To verify this, we stained brain sections from *Cntn1*^*-/-*^ (*Cntn1* KO) mice using the canonical PNN marker WFA. Unfortunately, *Cntn1* KO mice die within two to three weeks after birth, making it difficult to study PNNs in these animals since this is a relatively early stage in PNN development (39). However, we were able to utilize animals at postnatal day 16 (PND16), which is a timepoint in which immature PNNs are beginning to form. WFA staining revealed immature PNN staining in the *Cntn1*^*+/-*^ *(Cntn1* Het) brains comparable to PNNs observed in WT brains. Strikingly, WFA staining appeared aggregated and punctate on the surface of neurons from *Cntn1* KO animals even at this early PNN developmental stage (Fig. 6A). Our peak-node analysis revealed significant differences between *Cntn1* Het and *Cntn1* KO animals. Mean prominence or isolation of PNN staining peaks was significantly higher in *Cntn1* KO animals (Fig. 6B, mean prominence *Cntn1* Het = 1.99 ± 0.48 and *Cntn1* KO = 3.68 ± 1.37, *p* < 0.0001, two-tailed Student’s t test).

**Figure 6.**
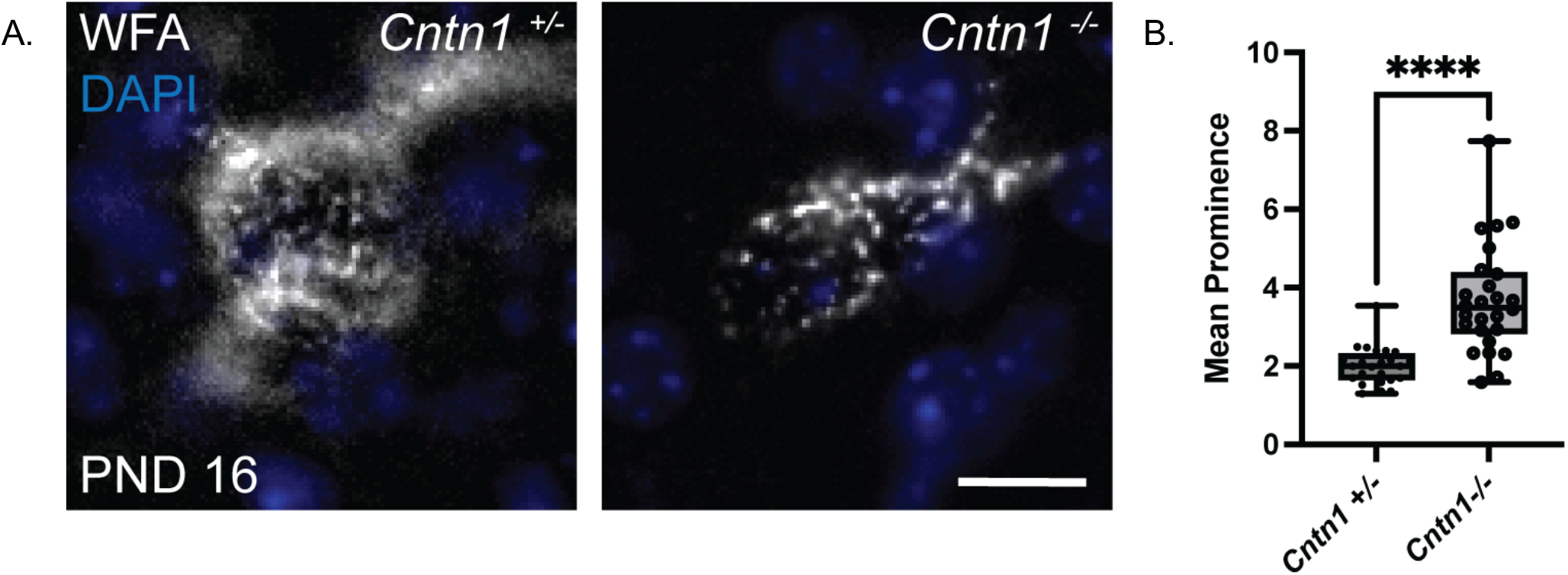
PNN structure is disrupted *in vivo* in mice carrying null alleles for *Cntn1*. *(A)* In mice, PNNs first appear around the second post-natal week of life, presenting as diffuse structures around subsets of cells in the CNS. At PND 16, PNNs in *Cntn1* ^+/-^ animals cortices show typical developing WFA-positive PNN staining around subsets of interneuron. In contrast in *Cntn1* ^-/-^ cortices PNNs form but even at this early developmental time point appear disrupted. WFA staining in these brains is aggregated and punctate compared to the PNN staining seen in *Cntn1* ^+/-^ animals. Structural analyses using *m*ean prominence or isolation of PNN peaks in these two groups was quantified using PNN node/ peak analysis. There were significant differences in the mean prominence of PNNs between the two groups of animals. Mean prominence index was significantly higher in *Cntn1* ^-/-^ animals (*Cntn1* ^-/-^ = 3.68 ± 1.37, *Cntn1* ^+/-^ = 1.99 ± 0.48 and p < 0.0001, two-tailed Student’s t test).

## DISCUSSION

An incomplete understanding of PNN structure and consequently our inability to manipulate them specifically, and in a cell-autonomous manner, has been a major hurdle towards understanding their function in the CNS. The vast majority of PNN components that have been identified are secreted molecules, and most are broadly expressed in the CNS (2, 14, 16, 19, 20, 28, 40–48). Yet, PNNs are conspicuous structures that form around a small and discrete population of cells, parvalbumin-positive interneurons. The cell-surface receptors responsible for PNN structure and their formation and around specific subpopulation of neurons has remained unclear. In our previous studies we identified two distinct interactions required for the proper formation of PNNs and demonstrated that PNN components bind to the neuronal surface by interacting with a HA backbone and through a complex formation involving Tnr and a soluble form of RPTPζ (22, 24). Here we focused on determining how the RPTPζ/Tnr complex attaches to the neuronal cell surface in PNNs. We believe the identification of the key receptor for this complex, Cntn1, represents a major step forward in understand PNN structure and ultimately will provide a key insight into understanding PNN function. In this current study, we utilized biochemical analyses and cell biological studies to first demonstrate that the RPTPζ/Tnr complex is attached via a GPI-linked protein. In *Ptprz1* KO neuronal cultures, Tnr is lost from the cell surface which made us hypothesize Tnr is anchored to the surface by RPTPζ and therefore, we focused our work on identifying a cell surface receptor for RPTPζ (22). Previous work had identified interactions between RPTPζ and the GPI-linked protein Cntn1 (35) and provided a structural basis for them (36, 37). Subsequently using a combination of PIPLC, function blocking antibodies and a RPTPζ protein variant that does not bind Cntn1, we showed that the interaction of RPTPζ with Cntn1 is indeed required for PNN structure. Finally, we demonstrated that PNN structure is disrupted *in vivo* as well in Cntn1 deficient animals. Furthermore, the disrupted aspect of PNN structure in *Cntn1* KO mouse brains phenocopies the *Tnr* and *Ptprz1* KO PNN phenotypes. Together our work demonstrates that Cntn1 is critical in binding RPTPζ and RPTPζ/Tnr complexes in PNNs to the neuronal surface and that Cntn1 is critical for PNN structure. We believe this work is the first to identify a key cell surface receptor for PNNs.

This work combined with our previous studies enabled us to create an updated model of PNN structure that we believe represent an important step forward to understanding PNN structures and function (Fig. 7). Importantly key features of this model were validated in our current study. For example, we had previously shown that aggrecan is bound in PNNs in a HA- and Ca^2+^-dependent manner (22). In contrast here we showed that despite the loss of aggrecan, Tnr remains bound to PNNs after HA digestion and Ca^2+^ chelation. Importantly, however, the combination of HA digestion and treatment with PIPLC eliminates both aggrecan and Tnr staining in PNNs. These results are predicted by our model and help highlight both the validity of the model and utility of using this model to direct future work on PNNs.

**Figure 7.**
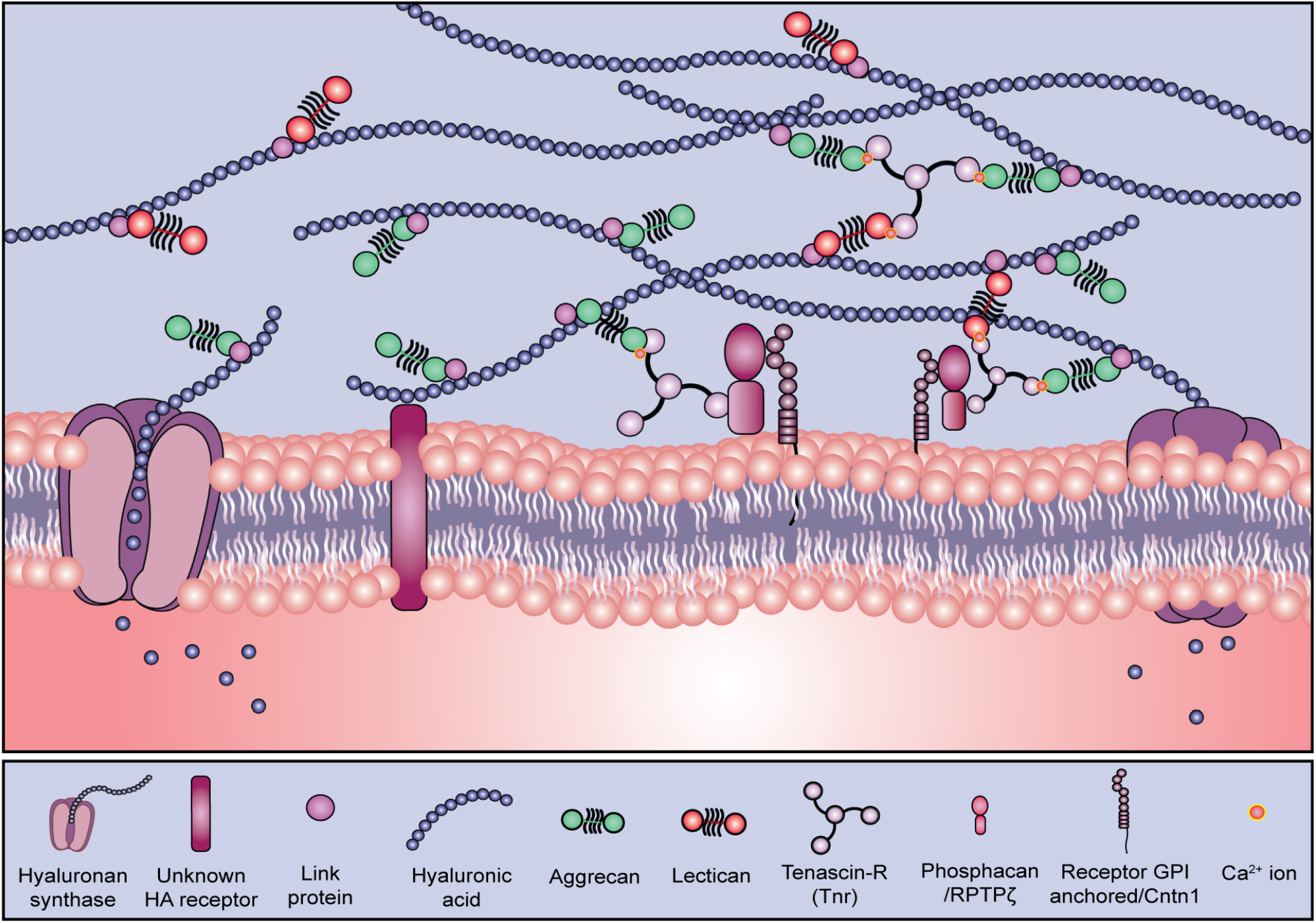
Proposed model of PNN structure. We propose a model for PNNs, in which lecticans are bound to the HA backbone and crosslinked to RPTPζ by Tnr. Our data suggests that it is the soluble form of RPTPζ, also known as phosphacan that mediates binding of PNN components to the cell surface. We present evidence that Cntn1 acts as a receptor for phosphacan in PNNs. To our knowledge this is the first direct demonstration of a PNN receptor. The mechanism by which HA binds to the cell surface is unknown and remains a goal of future studies.

Our data demonstrate that Cntn1 plays an essential role in immobilizing PNNs to the cell surface. However, it was a surprise to us that Cntn1 plays this critical role in PNN structure given that Cntn1 is a broadly and highly expressed during neural development (39, 49). In other words, Cntn1 is by no means uniquely expressed in PNNs. Furthermore, Cntn1 plays a number of key roles in neural development (36, 37, 45, 50–52). It is well known to play roles in axon growth and guidance, process fasciculation, oligodendrocyte development and myelination and synapse formation. The critical role Cntn1 plays in other neural development processes is highlighted by the fact that *Cntn1* KO mice are severely runted in postnatal development and live to a maximum age of 16-18 days due to failure of peripheral nerve innervation (50). In contrast, PNNs form on very selective and limited subset of interneurons (41). Although it is possible a specific splice variant of Cntn1 is expressed in PNN-bearing neurons, we do not believe Cntn1 is likely contributing to the cell-specificity of PNN formation *per se* but just to their structure. In support of this hypothesis, neurons that typically would have nets still stain positive for PNN components even in the *Ptprz1, Tnr*, and *Cntn1* knockouts, but PNN structure is disrupted from the typical reticular pattern to a more aggregated and punctate formation in these animals (15, 20, 22, 40, 46–48). Thus, we hypothesize that while a ternary complex of RPTPζ, Tnr, and Cntn1 is essential for PNN structure, it is likely that a HA-dependent binding of PNNs to the cell surface that gives these structures their cell type specificity. This is an area we are currently actively investigating. In addition, because of the many functional roles of Cntn1, functional analysis of the impact of PNN disruption knockouts was not possible. This is a similar challenge with the *Ptprz1* KO and *Tnr* KO mice in which the expressed proteins function in PNNs but also in other systems making understanding their particular role in PNNs difficult or impossible with current models. We believe, however, Cntn1 offers a key advantage going forward as it is a cell surface protein instead of a secreted protein. While secreted proteins can act at a distance from their cellular source, Cntn1 presumably acts in a cell autonomous manner. Therefore, conditional disruption of Cntn1 in PNN-bearing neurons could provide a unique tool to assess PNN functions. Overall, we believe this work provides an important step forward in understanding PNN structure and new insight into assess PNN functions.

## EXPERIMENTAL PROCEDURES

### Animals

Mice lacking the *Ptprz1* gene (*Ptprz1* KO) were generated as previously described (53) and received from Dr. Sheila Harroch (Department of Neuroscience, Institute Pasteur, Paris, France). For neuronal cultures, in addition to *Ptprz1* KO mice, timed pregnant CD-1 wildtype mice were purchased from Charles River Laboratory (Wilmington, MA, USA). All experiments carried out followed the protocols approved by the Institutional Animal-Care and Use Committee of SUNY Upstate Medical University.

### Antibodies

Mouse anti-tenascin-R 619 (MAB1624) and mouse anti-aggrecan (MAB11304) were purchased from R&D systems. Rabbit anti-aggrecan (AB1031) was purchased from Millipore Sigma (Burlington, MA, USA). Function blocking goat anti-contactin1 (AF904) and anti-contactin4 (AF5495) were purchased from R&D systems. Fluorescein labeled WFA (Wisteria floribunda agglutinin) was purchased from Vector Laboratories Inc. (Burlingame, CA, USA).

### Preparation of Homogenates, soluble and insoluble fractions for biochemical release assay

Brain homogenates for release assay of PNN component aggrecan, were derived from postnatal day 45 (PND 45) *Ptprz1* WT brains. Tissue was homogenized in 150mM sodium chloride and 50mM Tris with EDTA-free protease inhibitor tablets (Roche, Indianapolis, IN, USA 1 tablet in 10mL buffer), in a Potter Elvehjem homogenizer. Homogenates were centrifuged at 8,000g for 10 min at 4 °C. The supernatant was then removed, the pellet washed once and then resuspended in 1 mL buffer. A Bradford (Bio-rad) assay was performed, and protein concentrations were adjusted to 2.5mg/mL. Samples were treated with 2 μL chondroitinase ABC (Sigma-Aldrich, Saint Louis, MO, USA) and/or 5 μL PIPLC (Thermo Fisher) per 500 μl of sample and/or 1mM EDTA for 8 hours. Samples were centrifuged again at 8,000g for 10 min at 4 °C to separate soluble release (R) fraction and insoluble pellet (P) fractions and subsequently prepared for Western blotting by adding sample loading buffer and heating to 95 °C for 5 minutes.

### SDS-PAGE and western blotting

Protein concentrations were determined by Bradford assay before gel electrophoresis. 6-15% gradient SDS-polyacrylamide gels were used and transferred to 0.45 μM nitrocellulose membranes. Western blotting was conducted as previously described (54). Briefly, blots were placed in blocking buffer composed of 5% milk in low salt TBST and then incubated in primary antibody overnight. Blots were then incubated in HRP-conjugated secondary antibodies (The Jackson Laboratory, Bar Harbor, ME, USA) and exposed using supersignal west pico or femto chemiluminescent substrate (Thermo-Fisher Scientific, Rockford, IL, USA). Blots were imaged using Premium X-Ray film (Phenix Research Products, Candler, NC, USA).

### Purification of phosphacan

Phosphacan was purified by anion exchange chromatography as previously described (22, 55). In brief, the soluble fraction, from PND 3-4 CD-1 mouse brain, was filtered using a PVDF 0.22 μM filter, brought to a 0.5 M NaCl concentration, and run through a 1 mL HiTrap-Q HP column using a peristaltic pump connected to an Amersham Pharmacia RediFrac fraction collector (GE Healthcare Life Science). Sample was eluted over a continuous gradient of 0.5 M NaCl to 2.0 M NaCl over 10 column volumes and collected as 250 μL fractions. Fractions that were phosphacan rich, identified by dot blot analysis, were pooled, and concentrated using 100,000 MWCO Concentrators (AmiconUltra, EMD Millipore). Approximately 250 ng of purified phosphacan was added to *Ptprz1 KO* cultures after the first medium change at 3 DIV and 125 ng was added after the half-medium change at 6 DIV. Coverslips were fixed at 9 DIV and subsequently processed for immunocytochemistry.

### Preparation of RPTPζ-WT-Fc, RPTPζ-βdel-Fc

Early passage HEK293 cells were plated on 10cm dishes and transfected with RPTPζ-WT, RPTPζ-βdel constructs. Culture media was changed to Optimem, serum-free media 24 hrs post transfection. Media was collected 48 hrs post transfection and concentrated using 50,000 MWCO Concentrators (AmiconUltra, EMD Millipore). Protein concentration was estimated using a Bradford assay (Bio-rad).

### Primary Cortical Cultures

Neuronal primary cultures were prepared as previously described (19, 28). Briefly, cortices of embryonic day (E) 15-16 CD-1 WT or *Ptprz1* KO embryos were removed and digested in 0.25% trypsin-EDTA (ThermoFisher Scientific, Waltham, MA, USA). Mixed cells were filtered and suspended in Neurobasal medium with 3% B27, 1X Glutamax and 1X penicillin-streptomycin (ThermoFisher Scientific). Cells were then plated at a density of 2.1 × 10^6^ cells/mL on coverslips (500 μL/per well) pre-coated with poly-D-lysine (50μg/ml) and laminin (5 μg/ml) (Sigma-Aldrich, Saint Louis, MO, USA) in a 24-well dish. To remove glia, cells were treated with 5 μM cyotosine arabinoside (AraC, Sigma-Aldrich) at 1 DIV. The culture medium was then changed at 3 DIV to remove AraC and given a half change at 6 DIV. Cells were maintained at 37°C/ 5% CO_2_ until fixation. As per experimental requirements, coverslips were treated with 10 μL ChABC for 30 mins and/or 2.5 mM EGTA for 15mins and/or PIPLC for 30mins for biochemical release assay of PNN components in neurons. Coverslips were fixed using 4% paraformaldehyde (PFA) with 0.01% glutaraldehyde, pH 7.4 and subsequently processed for immunocytochemistry.

### PNN recovery in RPTPζ deficient cultures

Neuronal primary cultures were prepared as previously described from RPTPζ deficient, *Ptprz1* KO mice. Purified RPTPζ-WT or RPTPζ-βdel was added to cells at 2μg/ well of a 24 well dish at 3 DIV. Additional 1μg/ well was added after half media change at 6 DIV. Cells were fixed and analyzed at 9 DIV.

### PNN disruption in CD1 WT cultures

Neuronal primary cultures were prepared as previously described from CD1 WT mice (19, 22, 24, 28). Purified RPTPζ-βdel (2μg/ well of a 24 well dish) or RPTPζ-WT (2μg/ well of a 24 well dish) or anti-Cntn1 antibody or anti-Cntn4 antibody (2.5μg/ well of a 24 well dish) was added to cells at 6 DIV. Cells were fixed and analyzed at 9 DIV.

### Immunocytochemistry

Primary cortical cultures plated on coverslips were fixed at 9 DIV in cold 4% phosphate-buffered paraformaldehyde (PFA) with 0.01% glutaraldehyde, pH 7.4. Cells were then blocked in screening medium (DMEM, 5% FBS, 0.2% sodium azide) for 1 hour, before adding primary antibodies overnight at 4°C. The following day, Alexa-fluor conjugated secondary antibodies (ThermoFisher Scientific) in screening medium were added to the cells for 2 hours before mounting the coverslips with ProLong anti-fade kit (ThermoFisher Scientific). Cell nuclei were visualized with Hoechst solution (ThermoFisher Scientific) diluted in 1x PBS. Coverslips were imaged using an epi-fluorescent Zeiss Imager.A2 with Nikon Elements software package. Final images were gathered and formatted using Fiji software (56) and assembled into figures using Adobe Photoshop and Adobe Illustrator.

### Quantification and Statistical analyses

PNN component intensity was calculated by taking large scan images (1200μm x 900μm) from 3 different areas for each coverslip. A (64μm x 64μm) area devoid of PNN staining was used for background subtraction and intensity was calculated using the measure function of ImageJ. PNN peak or node analysis were used to quantitatively describe the PNN aggregation seen on the surface of neurons in cultures derived from *Ptprz1* KO mice and CD-1 wildtype mice. PNN images (40μm x 40μm) were processed using the local maxima function of ImageJ to identify peaks (“nodes”) of intense PNN staining. Once the nodes were identified, an *ad hoc* algorithm was used to measure the difference in intensity between the nodes and their surrounding space on the cell surface (mean node prominence) was calculated and plotted for each genotype. Differences were found significant at p < 0.05 (unpaired Student’s t-test or analysis of variance (ANOVA) with Tukey’s *post hoc* analyses as appropriate) using Graphpad Prism or RStudio statistical software.

## Authors contributions

R. T. M. and S. B. conceptualization; R. T. M. and S. B. data curation; A. S., R. T. M., and S. B. formal analysis; R. T. M. and S. B. funding acquisition; A. S., G. N., R. T. M., and S. B. investigation; A. S., G. N., R. T. M., and S. B. methodology; R. T. M. and S. B. project administration; R. T. M. and S. B. supervision; A. S., R. T. M. and S. B. writing–original draft; A. S., R. T. M. and S. B. writing–review and editing.

## Funding and additional information

Research reported in this manuscript was supported by the National Institute of General Medical Sciences under award number R01 GM143757 (R. T. M. and S. B.). The content is solely the responsibility of the authors and does not necessarily represent the official views of the National Institutes of Health.

## Abbreviations

ANOVA: analysis of variance
Chondroitinase ABC: ChABC
CNS: central nervous system
Cntn1: contactin-1
CP: critical period
CSPG: chondroitin sulfate proteoglycan
DAPI: 4′,6-diamidino-2-phenylindole
DIV: days in vitro
E: embryonic day
ECM: extracellular matrix
GPI: glycosylphosphatidylinositol
hyaluronan: HA
PIPLC: phosphatidylinositol-specific phospholipase-C
PNN: perineuronal net
PND, postnatal day, RPTPζ: protein tyrosine phosphatase receptor type Z
Tnr: tenascin-R

